# High-resolution ab initio reconstruction enables cryo-EM structure determination of small particles

**DOI:** 10.1101/2025.09.08.674935

**Authors:** Kookjoo Kim, Huan Li, Oliver B. Clarke

## Abstract

Despite recent advances in data acquisition and algorithmic development, applying single particle cryogenic electron microscopy (cryoEM) to small proteins (<50kDa) remains challenging, even when high quality data are available, in part due to the lack of reliable low-resolution structural features to inform initial alignments. Here we present a workflow which effectively bypasses this step, by obtaining initial particle orientations directly from heterogeneous ab initio reconstruction in CryoSPARC solely using data at high spatial frequencies. Applying this approach, we solve the structure of a previously intractable protein in a publicly available dataset, iPKAc (EMPIAR-10252), 39 kDa, resolved at an estimated resolution of 2.7 Å as well as a hemoglobin alpha-beta dimer (EMPIAR-10250) at 29kDa, resolved to an estimated resolution of 4 Å. We also show that the Aca2-RNA complex (37kDa, EMPIAR-11918) can be resolved by this approach directly from a blob-picked particle stack, in a single round of heterogeneous ab initio reconstruction followed by local refinement. The map of iPKAc is of sufficient quality to autobuild 325 of 356 residues present in the original crystal structure using Modelangelo, and ordered ATP and magnesium ions can clearly be resolved. The Hb-dimer has clear secondary structural features, identifiable hemes, and visible bulky sidechains, consistent with the estimated resolution. We expect that this approach may be useful for cryo-EM analysis of other small particles near or below the theoretical size limit.

## Introduction

The advent of high resolution cryo-EM over the past two decades has revolutionized structural biology^1^, allowing structural characterization of macromolecules that were previously intractable either due to compositional or compositional heterogeneity, or in general unwillingness to form ordered crystals. One remaining limitation of modern cryoEM methods is the size limit to which this method can be applied. Below ∼50kDa, obtaining an initial reconstruction with interpretable structural features becomes challenging or, in many cases, impossible with current approaches, with theoretical calculations suggesting a limit of around 38kDa^2^.

One solution to this problem involves the use of so-called “fiducials” - rigid binders that increase the effective ordered mass of the macromolecule, thereby facilitating assignment of initial orientations, after which high resolution refinement can converge^3^. However, fiducials suffer from their own limitations - identifying them is laborious and target specific, dissociation during sample preparation can be a problem, and even if a binder is identified, it may impinge on a functionally relevant part of the protein, such as the ligand binding site.

A second approach under consideration is the development of direct electron detectors with improved quantum efficiency at 100kV, which calculations suggest should produce particle images with increased signal-to-noise ratio for small macromolecules in thin ice for a given electron fluence^4^. This approach is very promising, but still under development, and not yet available for general use.

New computational approaches are therefore needed for direct cryo-EM characterization of small proteins. One such approach was recently described, involving use of a data driven regularization strategy (BLUSH) to facilitate convergence from an initially poor-quality map, generated by stochastic gradient descent at low resolution, to a high-resolution refinement^5^. Here, we describe a complementary approach, using a cryoSPARC-based workflow, that allows generation of a buildable map in a single step, using high resolution heterogeneous ab initio reconstruction (abbreviated HR-HAIR).

## Results

HR-HAIR facilitates high resolution reconstruction of small particles by cryo-EM, near the theoretical size limit, by restricting *ab initio* reconstruction to use data only at high spatial frequencies. While here we have implemented this protocol using CryoSPARC^6^, it is reasonable to expect the same approach to be possible to implement in RELION, which also has a stochastic gradient descent-based *ab initio* reconstruction module^7^.

The key step in this workflow is heterogeneous *ab initio* reconstruction, as implemented in CryoSPARC, but using a specific set of non-default parameters, namely using only a small window of data at high spatial frequencies for initial model generation and initial orientation assignment, and using a very small resolution step size (Fourier radius step) between iterations. This was motivated by the observation that iPKAc (EMPIAR-10252)^8^ shows very clear high-resolution features in 2D classification for multiple different orientations, but has so far not yielded an interpretable reconstruction using conventional approaches (**Fig 1a**). The specific parameters used for *ab initio* reconstruction and 2D classification for each dataset are detailed in the Methods. We find that by only using high spatial frequency information, and critically, by reducing the Fourier radius step between iterations, we are able to converge on an interpretable map for iPKAc (EMPIAR-10252) (**Fig 1 b**), where other approaches that we and others have tried produced maps with no interpretable high resolution structural features, despite the fact that high resolution features are clearly visible in 2D class averages (**Fig 1a**).

**Figure 1:**
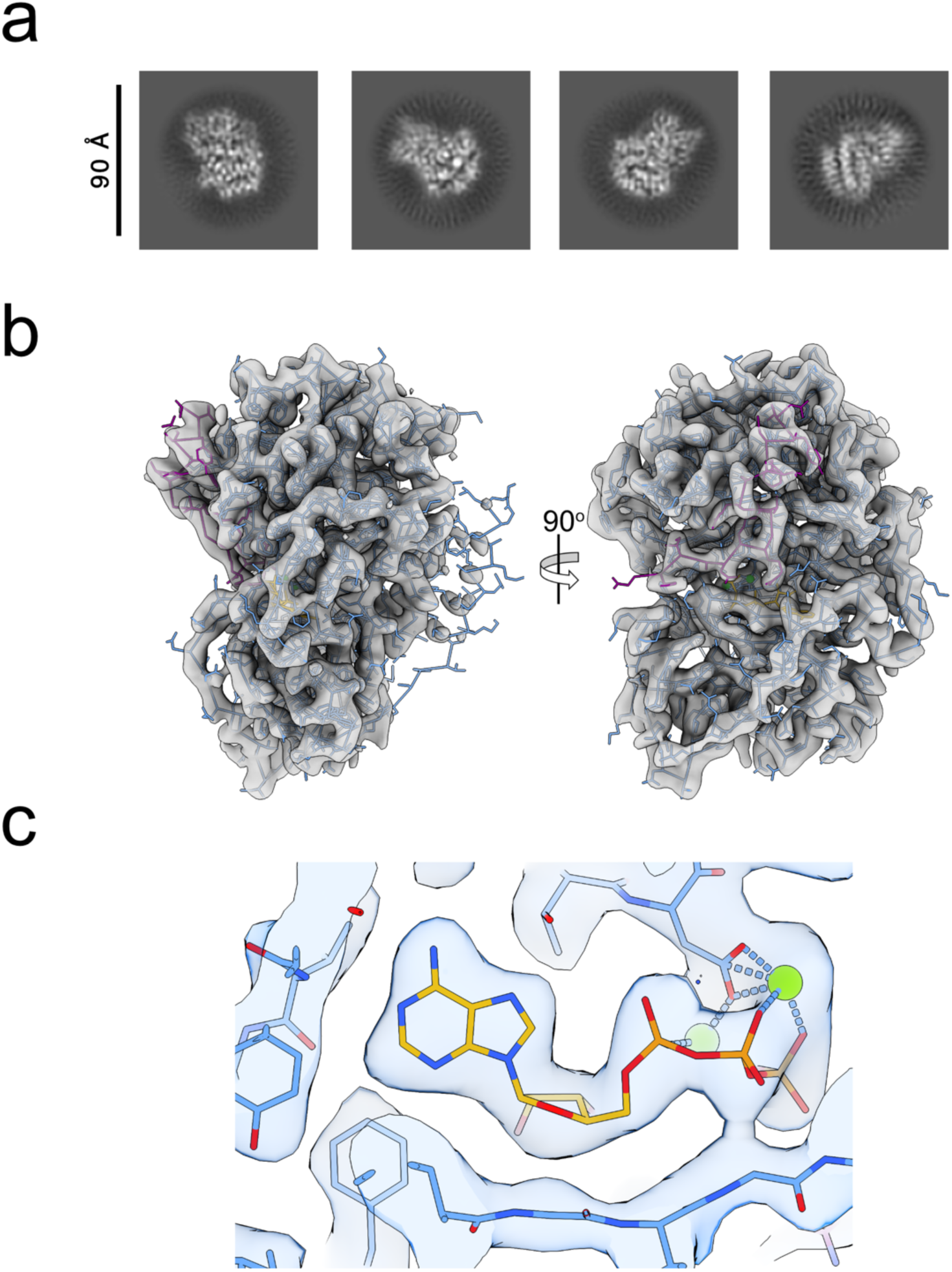
Structure of iPKAc solved using HR-HAIR. a) 2D classes of iPKAc (EMPIAR-10252; dataset 18mar03b) exhibit high resolution features in multiple orientations. Classes shown here have been cropped to exclude blank areas outside of spherical mask. b) Two views of the structure of iPKAc, fit to the unsharpened map after *ab initio* reconstruction and local refinement, represented as a C-alpha trace with sidechains shown. The catalytic domain of PKA is represented in light blue, the inhibitory peptide in purple, and the map as a transparent gray surface. The disordered N-terminal helix can be seen on the righthand side of the left view. c) Closeup of the sharpened map around the ATP binding site, with all atoms shown.

The map of iPKAc shows clear density for the kinase domain and inhibitory peptide, bound ATP and two resolved magnesium ions (**Fig 1 c & Movie S1 & S2**). These features are evident in the initial map obtained from ab initio orientations (**Fig S1**), and can be further improved by local refinement. Notably, the N-terminal helix is almost completely disordered in this structure (**Fig 1 b**), in contrast to the crystal structure^9^. The map is of sufficient quality to facilitate autobuilding of 325/356 residues using Modelangelo^10^ (**Fig S2 e**), demonstrating that the residual preferred orientation does not significantly impede structural interpretation. Notably, the map directly out of *ab initio*, prior to any further refinement of orientations, facilitated autobuilding of 264/356 residues, with clear density for the ATP and magnesium ions.

We also applied the same approach to the hemoglobin (Hb) alpha-beta heterodimer (EMPIAR-10250)^8^, which at 29.4kDa with pseudo-C2 symmetry represents an extremely challenging target for cryoEM analysis, near the limit of current capabilities. An additional complication of this dataset is that the Hb-dimer is a minor species in a population that is primarily composed of Hb tetramers. Similarly to iPKAc, high-resolution features were evident in initial 2D class averages (**Fig 2 a**), motivating us to attempt reconstruction. By applying the same workflow to the hemoglobin dimer (**Fig S3**), we were able to obtain a reconstruction at an estimated resolution of 4 Å (**Fig 2 b & Movie S3 & S4**), with clear sidechain density in the center of the map, clear secondary structure throughout, and interpretable density for the two heme ligands (**Fig 2 c**). As a measure of map quality and interpretability, we ran Modelangelo to autobuild the model based on the sequence; Modelangelo was able to build 163/286 residues (**Fig S4 e**), which is consistent with the estimated resolution of 4Å.

**Figure 2:**
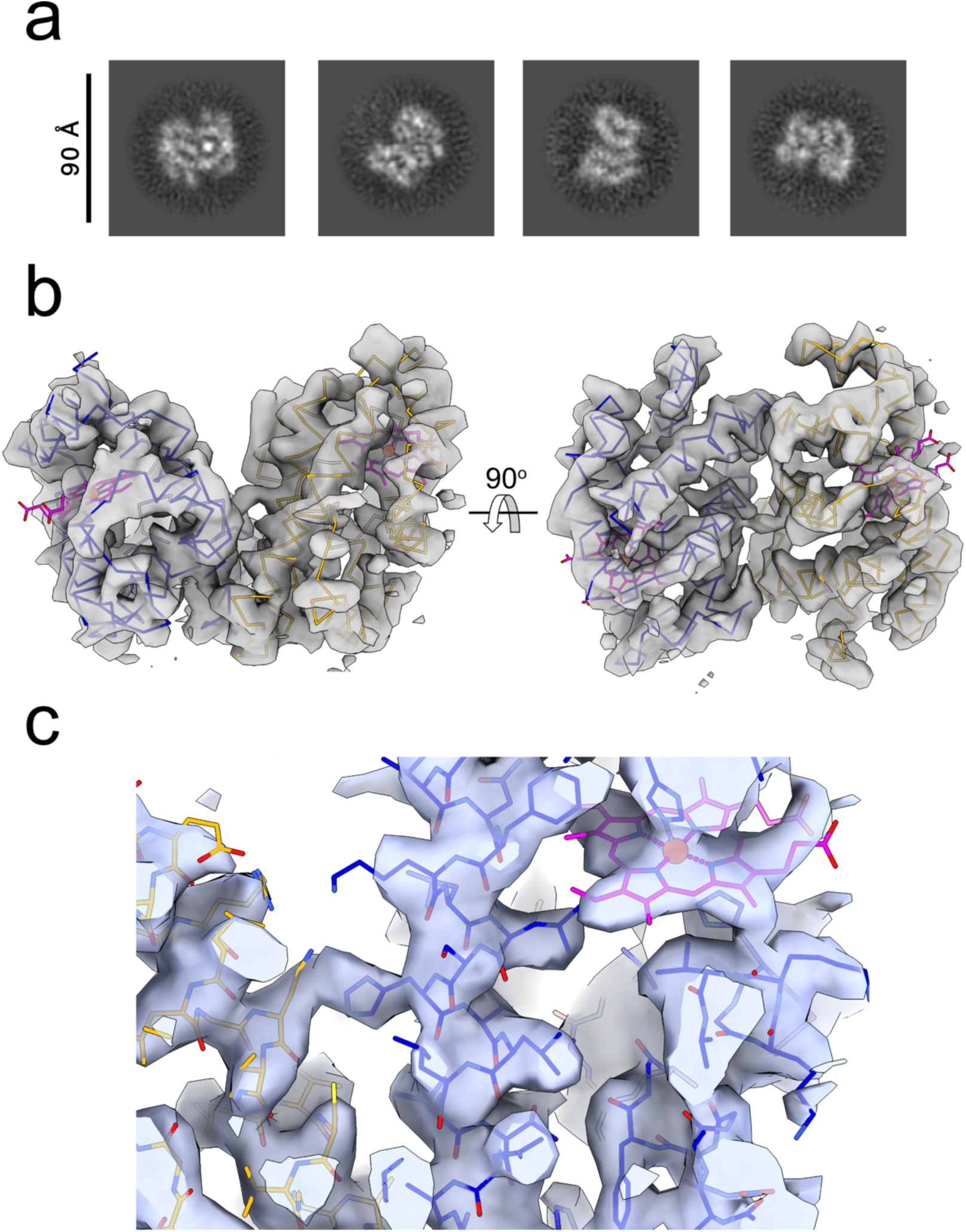
Structure of hemoglobin dimer solved using HR-HAIR. a) 2D classes of Hb-dimer (EMPIAR-10250; dataset 18feb21c) exhibit high resolution features in multiple orientations. Classes shown here have been cropped to exclude blank areas outside of spherical mask. b) Two views of the structure of Hb-dimer, fit to the unsharpened map after *ab initio* reconstruction and local refinement, represented as a C-alpha trace with hemes shown. The alpha chain is colored blue, the beta chain orange, and the hemes magenta. c) Closeup of the sharpened map around the near the heme binding site in the alpha chain, with all atoms shown.

We next attempted to tackle the 37kDa Aca2-RNA complex (EMPIAR-11918), which has been previously reported to require a data-driven regularization approach in both classification and refinement to obtain a high-resolution reconstruction^5^. Here, we were able to obtain a nominally 3.0 Å map directly from a single round of 3-class HR-HAIR (**Fig S5**), initialized directly from blob picked particles, with no prior 2D classification (**Fig 3 b**). In a parallel approach, we performed single-class high resolution ab initio reconstruction followed by local refinement after a single round of 2D classification, resulting in a nominally 2.8Å map without further classification (**Fig 3 a & Movie S5 & S6**), which facilitated autobuilding of 196/234 protein residues using Modelangelo (**Fig S6 e**). The HR-HAIR approach, combining high resolution search with a very small resolution step between iterations, reproducibly converged upon the correct solution, whereas other parameters often converged on false minima that could not be successfully refined. The final resolution for the Aca2-RNA structure was 2.8 Å, which is somewhat lower than the published resolution of 2.6 Å. However, we obtained this from processing only the first 1245 micrographs of the 11232-micrograph dataset, and did not attempt a second round of HR-HAIR, so further improvements by the same approach appear achievable.

**Figure 3:**
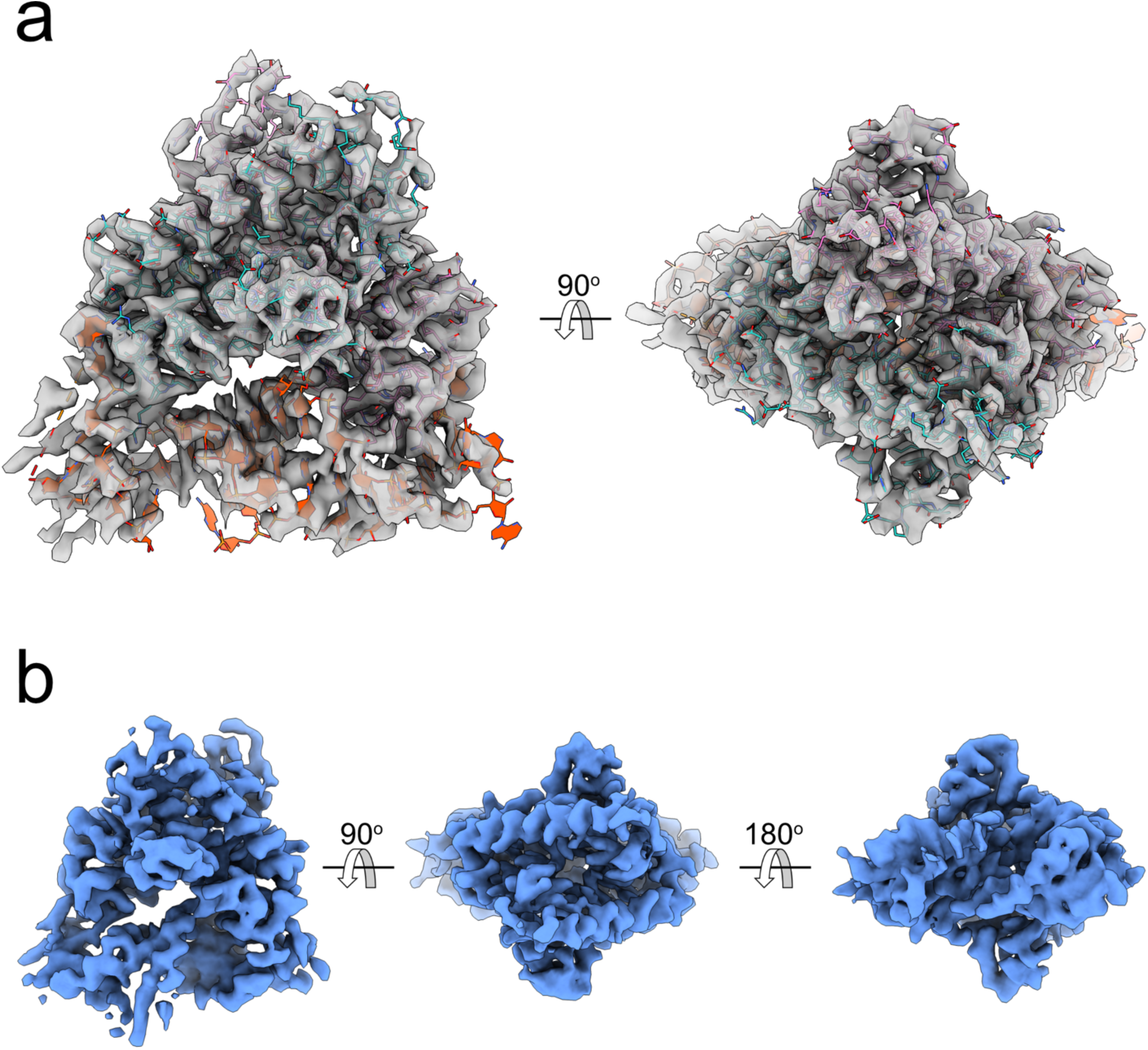
Structure of Aca2-RNA (EMPIAR-11918) solved using HR-HAIR. a) Sharpened map obtained after 1 round of 2D classification, and a single class HR-*ab initio* followed by local refinement, with atomic model (8W35) fitted. b) Three views of the unsharpened map obtained after a single round of 3-class HR-HAIR starting from the unclassified, blob-picked stack.

In all three cases, non-uniform or homogeneous refinement using the particle subset identified by HR-HAIR failed to further improve map quality, using initial lowpass resolution cutoffs up to 6 Å, but gave structural features consistent with the maps produced by local refinement of *ab initio* orientations (with a somewhat higher degree of anisotropy), suggesting that the initial alignments calculated in the *ab initio* step are sufficiently accurate to only require local searches for optimization.

The same approach applied to an even smaller protein, calmodulin (16kDa), produced recognizable bilobal features in 2D class averages (**Fig 4 a**), and domain level features in 3D (**Fig 4 b & Movie S7**), but no clear secondary structural features. However, the fact that structural features can be recognized for a protein of this size, using a relatively small particle stack and initial dataset, suggests that with a larger dataset, improved detectors, or lower accelerating voltage, structure determination of proteins below 20kDa using cryo-EM may be within reach.

**Figure 4:**
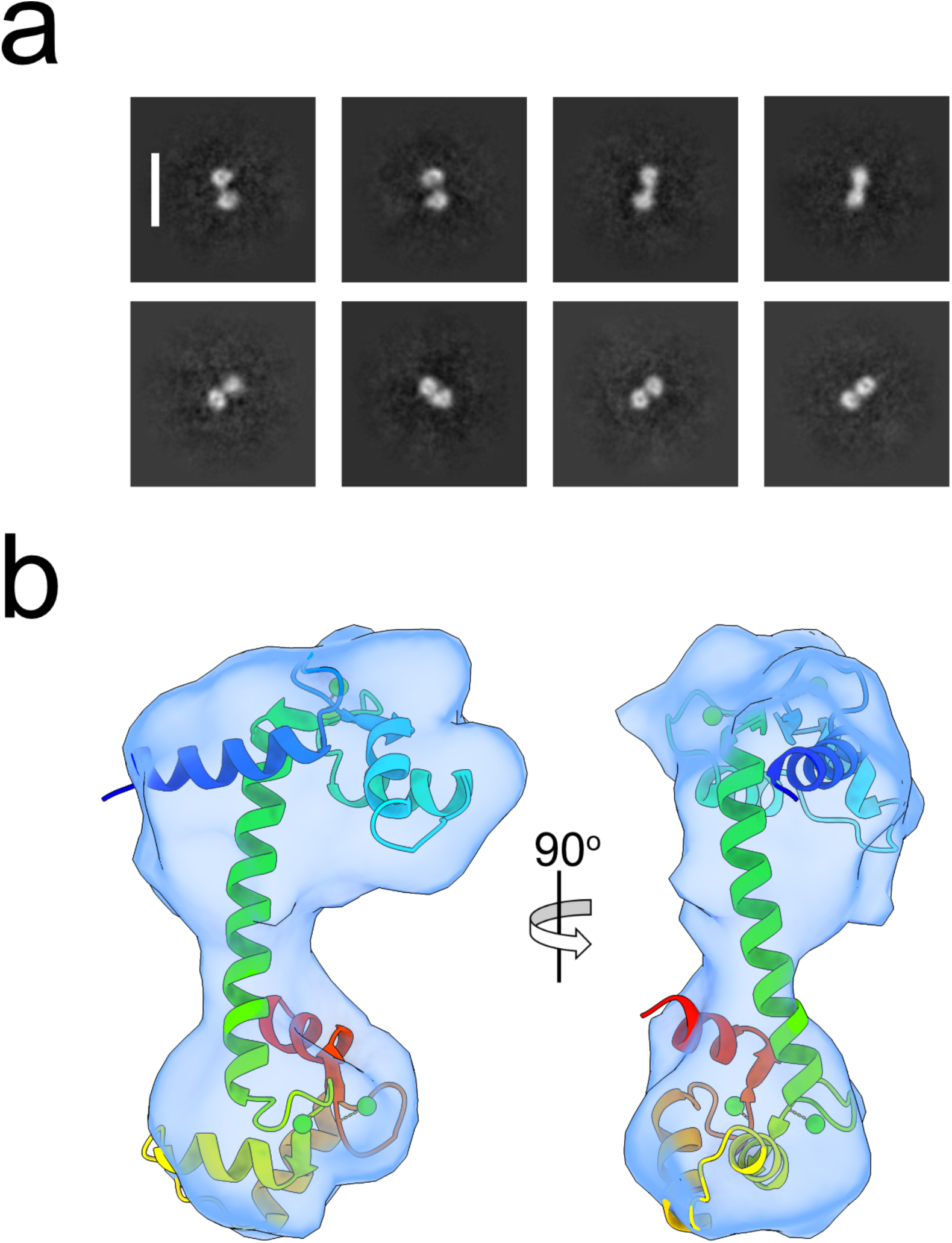
Domain-level features in a sub-20kDa protein by cryo-EM. a) Selected 2D classes of calmodulin. The white scale bar is 110 Å. b) *Ab initio* reconstruction of calmodulin, with structure 1CLL rigid fit to the density map.

We have not attempted using regularization functions other than non-uniform regularization during local refinement of the initial orientations. It is likely that application of BLUSH regularization or other data-driven regularization approaches either during local refinement or during the initial phase of high-resolution *ab initio* reconstruction may further improve upon the results reported here.

As has been noted^11^, run times increase with resolution for stochastic gradient descent, making this approach somewhat slower than using conventional *ab initio* parameter choices. For example, the 3-class HR-HAIR run for iPKAc took 9h30min on a single A6000 GPU. However, in the examples discussed here, a single round of *ab initio* accomplished a result which we were not able to obtain by other approaches (iPKAc and Hb-dimer), or which previously required multiple rounds of classification and refinement (Aca2-RNA).

In order to facilitate reproduction of these results, and easy application of other algorithmic approaches and methods development for the same particle stacks, upon publication we will deposit to EMPIAR the raw particle stack, the 2D-classified stack, and the final stack used for high resolution refinement, as well as the maps, masks and atomic models, which will be deposited to the EMDB and PDB respectively.

## Discussion

Stochastic gradient descent, the algorithm used for *ab initio* reconstruction in both RELION and cryoSPARC, is conventionally used to obtain an initial low-resolution volume, which is then used, after discarding particle alignments, as a reference for iterative assignment and refinement of particle orientations.

Here we show that by using data only at high spatial frequencies, it is possible to use stochastic gradient descent to obtain a map of sufficient quality for model building directly, which can be further improved by local refinement of particle orientations.

The fact that *ab initio* reconstruction alone is able to generate interpretable maps for iPKAc and Aca2-RNA, even in the presence of significant preferred orientation, suggests that the use of this approach for high resolution reconstruction and classification may be currently underestimated; further developments, including the addition of regularization functions and a gold-standard split, could potentially further improve the applicability of this approach for high-resolution structural studies. In particular, this is further emphasized by the fact that this approach generated an interpretable map for Aca2-RNA directly out of ab initio starting from a picked, unclassified stack, without prior 2D or 3D classification.

The iPKAc particle orientation distribution shows signs of what would usually be considered a severe preferred orientation, but the map is quite readily interpretable, with few signs of significant anisotropy. This fits with previous data showing that the preferred orientation problem, as such is less a problem of missing orientations, and more a problem of misalignment – particles from the preferred orientation being assigned to regions of Fourier space that are unoccupied^12,13,14^. It is possible that by only considering high resolution spatial information, the workflow described here suffers less severely from misalignment, in a complementary manner to data-driven regularization approaches towards the same goal^5^.

A 4Å map, with clear evidence for sidechains and bound hemes, was obtained for the 29kDa hemoglobin heterodimer, from a 13k particle stack originating from 588 movies collected on a K2 detector, with a Talos Arctica. The fact that an interpretable map could be obtained from this limited dataset for a sub-30kDa protein suggests that the estimated size limit for single-particle cryo-EM may need to be re-evaluated.

In CryoSPARC, *ab initio* reconstruction is performed without splitting the dataset into half-sets. Because, in the approach described here, the particle orientation information is retained after *ab initio*, the reported resolutions after local refinement cannot be assumed to be accurate. However, in the cases described here, the structural features of the map visually match the reported resolution. For example, in the case of iPKAc, the reported resolution is 2.7 Å, and magnesium ions, carbonyl bumps and small sidechains can clearly be resolved, and successful Modelangelo autobuilding was achieved. Furthermore, performing non-uniform refinement using the map generated by HR-HAIR, but with particle orientations discarded, gives reported resolution numbers which are comparable, albeit with a visually slightly inferior map.

Another limitation of this approach is that this is only likely to succeed for small particles that are relatively rigid. For small particles that show some degree of continuous flexibility, this is unlikely to be a viable approach due to the absence of consistent high resolution structural features. Nevertheless, a significant group of small kinases and other enzymes fall into the former category, for which structure determination at high resolution can inform structure-based drug design, suggesting that this approach may be of some utility for proteins that are currently challenging to resolve.

## Data availability

All maps, models, masks and particle stacks will be available at the EMDB, PDB and EMPIAR upon publication. Movies and particles for calmodulin will be deposited to EMPIAR upon publication. In the meantime, preliminary maps and models can be downloaded here:

https://www.dropbox.com/scl/fi/l8br619vb0mqx3aqvh64e/maps_and_models_for_ms.zip?rlkey=nh9428hm5e00uxoxmltvy1rlu&dl=0

Supplementary Movies can be downloaded here:

https://www.dropbox.com/scl/fi/tj11u726pe1kcp7qvx0b7/supp_movies.zip?rlkey=5jqjoam0hl6wyi45ib0r94dil&dl=0

## Supporting information

Movie S1 (iPKAc overall)

Movie S2 (iPKAc closeup)

Movie S3 (Hb-dimer overall)

Movie S4 (Hb-dimer closeup)

Movie S5 (Aca2-RNA overall)

Movie S6 (Aca2-RNA closeup)

Movie S7 (CaM map/model)

## Acknowledgments

Cryo-EM data for calmodulin were collected at the Columbia Cryo-EM center and at the Simons Electron Microscopy Center (SEMC), with the assistance of staff from both SEMC and the Columbia University Cryo-EM Microscopy Center. R. Grassucci and Z. Zhang from the Columbia Cryo-EM Center assisted with data collection. We thank all the authors who deposited their data to EMPIAR, making this study and many others possible.

## Author contributions

O.B.C. conceived the study. K.K. designed and executed the experiments with calmodulin, and H.L. and K.K. prepared figures. All authors analyzed the results. O.B.C wrote the manuscript in consultation with all authors.

## Methods

### Image processing

All operations were performed in CryoSPARC v4.7.1 for Aca2-RNA, and using a beta version of CryoSPARC v5, which has an option to apply a soft spherical mask to the ab initio volume at each iteration, for iPKAc and the Hb-dimer. All datasets were initially preprocessed using Patch Motion, Patch CTF and Micrograph Denoiser (trained on each individual dataset; see **Fig S7**), and picking was performed on the denoised micrographs (with extraction from the non-denoised, dose-weighted averages).

### Image processing of iPKAc

Image processing of iPKAc was initialized from the 18mar03b dataset (1353 movies), part of the EMPIAR-10252 entry. Blob picking on denoised micrographs using elliptical templates (50-70 Å) gave 854k initial picks. A round of template picking (performed using 20Å filtered projections of an initial, anisotropic consensus refinement) identified 787k particles, which were extracted in an initial box of 384px (0.5585 Å/pixel) and binned to 192 px (1.117 Å/pixel). This particle stack was used for all subsequent processing unless otherwise indicated. 2D classification was performed with the following custom parameters:

*200 classes, maximum reconstruction resolution of 3 Å, initial classification uncertainty factor of 1, Circular mask diameter 80 Å, number of O-EM iterations 80, number of final full iterations 20, Batchsize per class 400*.

Classes with well-defined high-resolution features – clear helicity and bulky sidechains – were chosen for subsequent high resolution ab initio processing, totaling 154k particles.

One round of HR-HAIR was performed using the 154k particle stack as input to *ab initio* reconstruction, requesting three classes and the following custom parameters (run time 9h 30m, single A6000 GPU):

*Initial resolution 5 Å, maximum resolution 2.3 Å, center structures in real space OFF, Fourier radius step 0.005, initial minibatch size 300, final minibatch size 1000*.

Of the resulting classes, one class (53k particles) had well-defined high-resolution features (sidechains, ligands and helices), while the other two were not interpretable.

The good class was reconstructed after re-extraction without binning (0.5585 Å per pixel, 384 px box size) by homogeneous reconstruction, redoing the gold-standard split, and locally refined with a rotation search extent of 2 deg, shift search extent of 1 Å, recentering rotations and shifts at each iteration, with an initial lowpass resolution of 6 Å. This generated the final map, with an estimated resolution of 2.7 Å.

### Image processing of Hb-dimer

Image processing of the Hb-dimer was initialized from the 18feb21c dataset (588 movies), part of the EMPIAR-10250 entry. Blob picking on denoised micrographs using elliptical templates (40-60 Å) gave 233k initial picks, which were extracted in an initial box of 384 px (0.5585 Å per pixel) and Fourier cropped to 192 px (1.117 Å per pixel). 2D classification was performed with the same custom parameters as for iPKAc, and classes corresponding to the dimeric species with high-resolution features were selected, totaling 21k particles. A 15k subset of these particles (selected based on curation of the most promising micrographs) was used to train a Topaz model which was used to repick the dataset, resulting in 48k particle stack after 2D classification. This stack was used as the starting point for subsequent processing. Two rounds of HS-HAIR requesting two classes were performed with the following custom parameters:

*Initial resolution 5 Å, final resolution 3 Å, Fourier radius step 0.005, window inner/outer diameter 0.4/0.5 (CryoSPARC v5-beta specific), center structures in real space OFF, enforce non-negativity FALSE, initial minibatch size 300, final minibatch size 1000*.

The good class was used to initialize the second round of HS-HAIR. The final class from the second round included 14k particles. After homogenous reconstruction, the nominal resolution was 3 Å. Local refinement using the same parameters as for iPKAc resulted in a final map with a nominal resolution of 3.2 Å. Non-uniform refinement with an initial lowpass resolution of 8 Å gave a reconstruction with a nominal resolution of 4Å; this is reported as the estimated resolution, due to uncertainty about the true resolution of the maps derived from *ab initio*.

### Image processing of Aca2-RNA

Image processing of Aca2-RNA was initialized from the first 1245 movies of the EMPIAR-11918 entry. Patch Motion in this case was performed with an output F-crop of 1/2, resulting in a final pixel size for the dose-weighted averages of 1.06 Å. Blob picking on denoised micrographs using elliptical templates (50-75 Å) gave 743k particles, which were extracted in a box-size of 256 px. The raw, unclassified stack was used for a test run of HS-HAIR, requesting three classes, with the following custom parameters:

*Initial resolution 5Å, final resolution 3 Å, Fourier radius step 0.005, center structures in real space OFF, enforce non-negativity FALSE, initial minibatch size 300, final minibatch size 1000*.

This resulted in a single good class, with clear helical protein and nucleic acid density, and two uninterpretable, anisotropic classes. The good class (254k particles) was used for subsequent local refinement, resulting in a map with a nominal resolution of 3.0 Å.

2D-classification of the unclassified stack was performed with the same custom parameters as for the other two datasets. Good classes comprising 219k particles were selected for a single class *ab initio* reconstruction with the same custom parameters as outlined above.

The particles and the volume from the single-class *ab initio* were locally refined with a rotation search extent of 2 deg, shift search extent of 2 Å, recentering rotations and shifts at each iteration, with an initial lowpass resolution of 6 Å. This generated the final map, with an estimated resolution of 2.8 Å.

### Data collection and image processing of calmodulin

Recombinant His-tagged calmodulin was expressed and prepared as previously described^15^. Grids were prepared by applying 3µL purified calcium-bound calmodulin (10 mg/mL) to 0.6/1 µm UltrAuFoil grids (Quantifoil/SPT LabTech), blotting in a Vitrobot Mark IV (15s blot time, 30s wait time), and vitrifying in liquid ethane. 306 movies were collected on a Titan Krios operated at 22,500X magnification (pixel size 1.06 Å), equipped with a Gatan K2 direct electron detector. After Patch Motion and Patch CTF, micrographs were denoised using Topaz Denoise^16^, and particles were picked manually and used to train a Topaz neural network model, which resulted in the extraction of 80k initial particles, extracted in a box size of 256px (1.06 Å/pixel). 2D classification was performed with the same custom parameters as for the other three datasets, except that the maximum reconstruction resolution was set to 6 Å. 2-class HR-HAIR was performed on the initially picked stack, with the following custom parameters:

*Initial resolution 6Å, final resolution 5 Å, Fourier radius step 0.005, center structures in real space OFF, enforce non-negativity TRUE, initial minibatch size 300, final minibatch size 1000*.

This resulted in two classes, one of which had a distinct dumbbell-like bilobal shape consistent in dimensions with calmodulin.

### Model fitting and refinement

In the case of iPKAc, the crystal structure (1ATP) of the peptide-inhibited PKA catalytic domain was oriented and rigid-body fitted in the map using Chimera. Both rigid body fit and real space refined (using phenix.real_space refine^17^) models are provided. In the case of the Hb-dimer, the alpha and beta chains, with associated hemes, were extracted from the previous cryo-EM structure (6NBD) and rigid-body fitted. Both rigid body fit and real space refined (using phenix.real_space refine) models are provided For Aca2-RNA, the published cryo-EM structure (8W35) was fit directly to the map; Both rigid body fit and real space refined (using phenix.real_space refine) models are provided. As a method to quantify map quality and interpretability, we ran Modelangelo, as implemented in Relion 5, providing FASTA format sequences of the relevant components and the sharpened map.

## Figure generation

Figures were generated using UCSF ChimeraX^18–20^.

---

**Figure S1:**
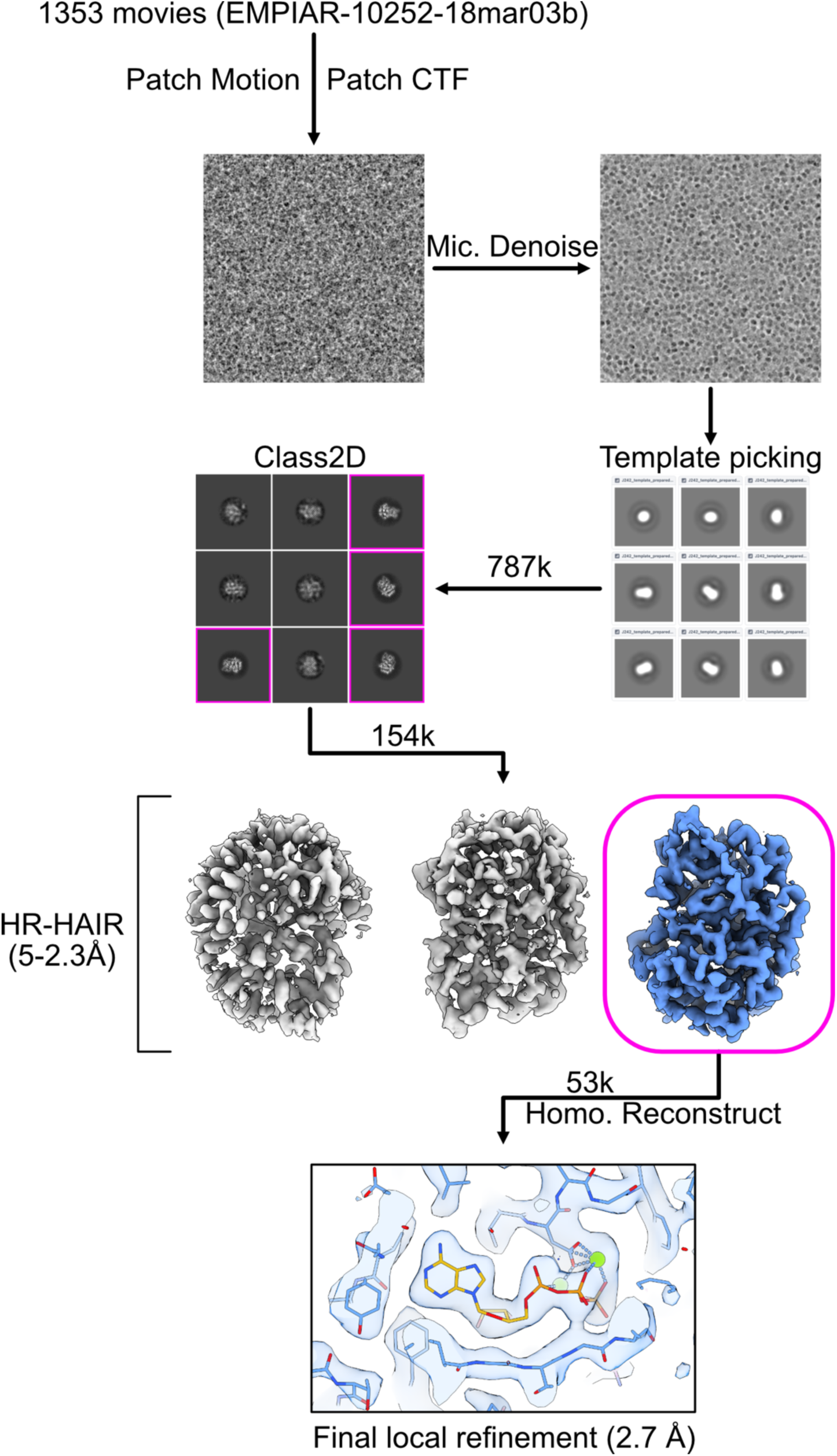
iPKAc image processing schematic. Image processing workflow overview for iPKAC; see Methods for details.

**Figure S2:**
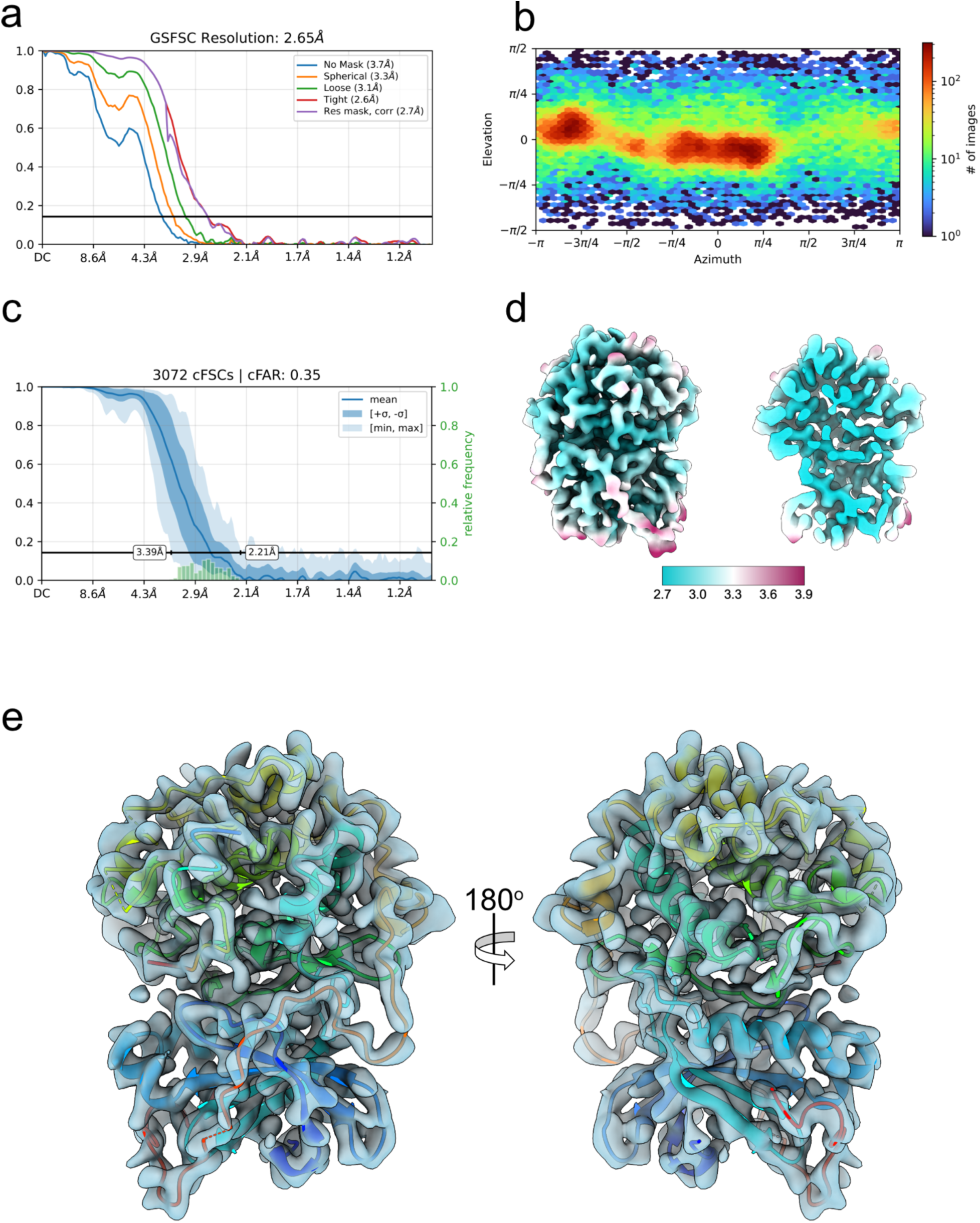
iPKAc FSC curves, local resolution and orientation distribution. a) FSC curves after local refinement with a 6Å initial lowpass filter. b) Particle orientation distribution. c) Conical FSC distribution and conical FSC area ratio. d) Unsharpened map colored by local resolution, calculated by the local resolution module of CryoSPARC, with an FSC=0.5 threshold. On the left, an overall view of the map; on the right, the same view, sliced down the middle in the plane of the page. e) Unsharpened map (transparent surface) with model autobuilt by Modelangelo (ribbons, spectral coloring)

**Figure S3:**
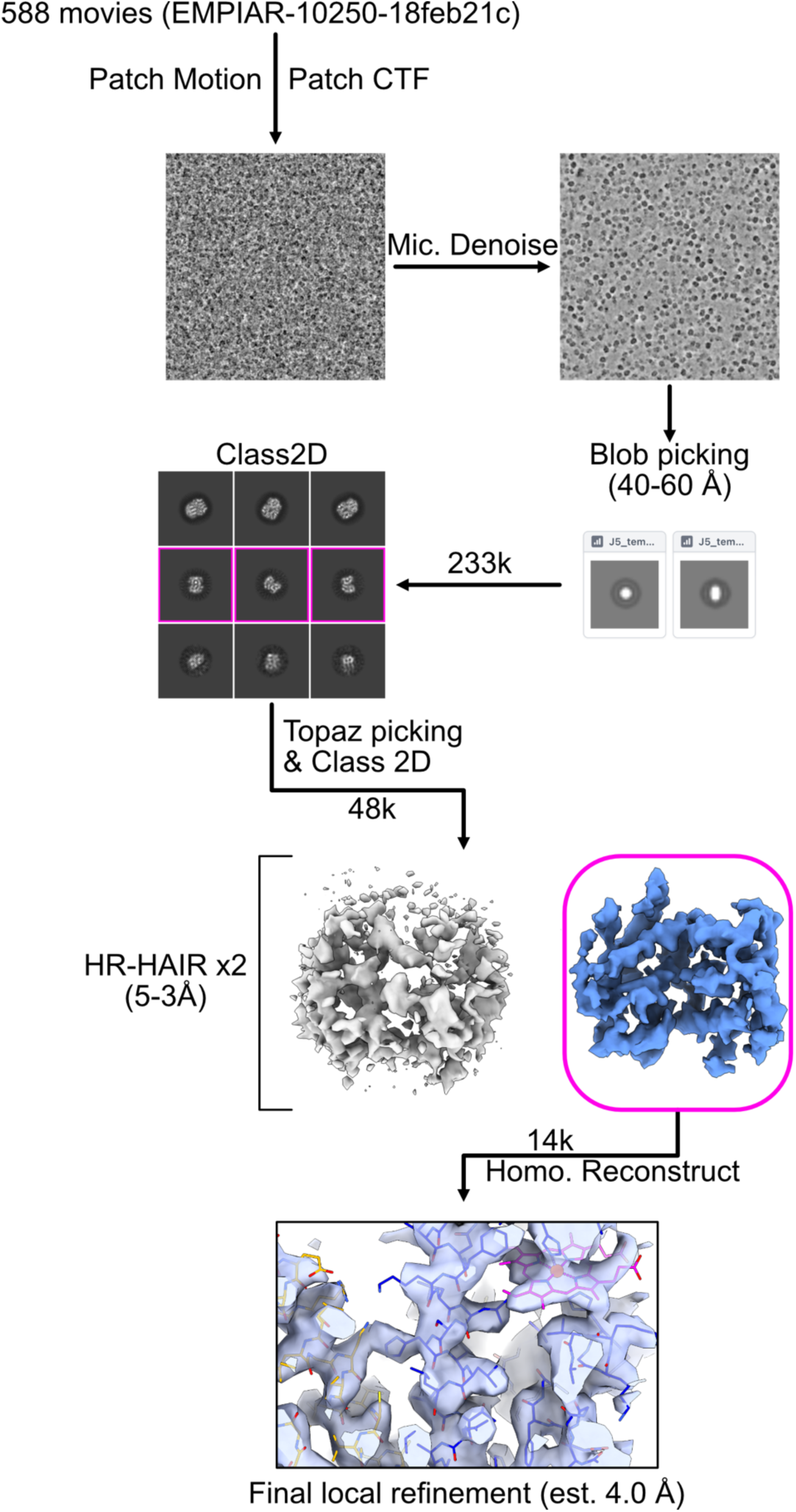
Hb-dimer image processing schematic. Image processing workflow overview for Hb-dimer; see Methods for details.

**Figure S4:**
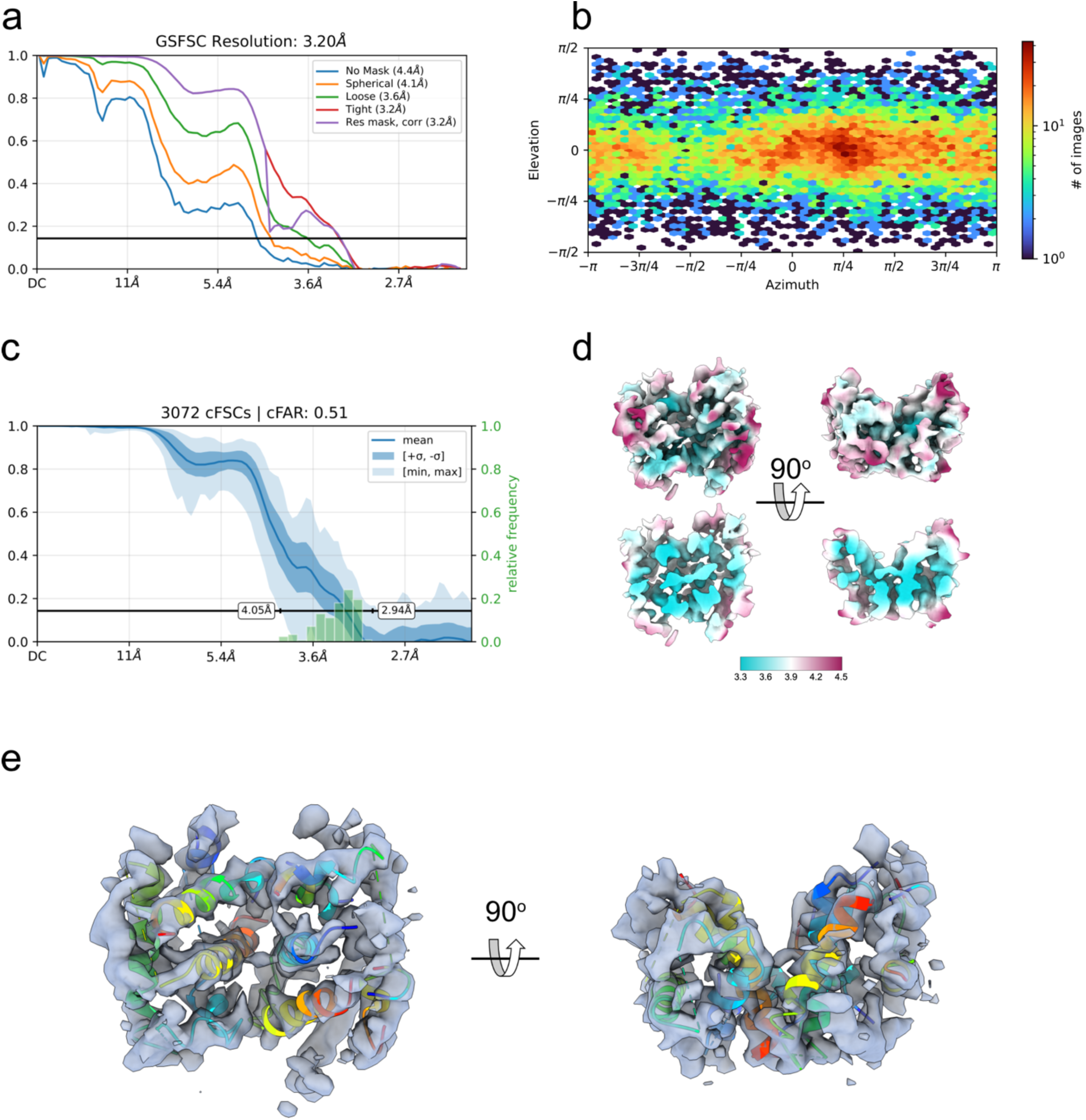
Hb-dimer FSC curves, orientation distribution and local resolution. a) FSC curves after local refinement. b) Particle orientation distribution. c) Conical FSC distribution and conical FSC area ratio. d) Two views of the unsharpened map colored by local resolution, calculated by the local resolution module of CryoSPARC, with an FSC=0.5 threshold. Upper panels show an overall view of the map; below, the same view, sliced down the middle in the plane of the page. e) Unsharpened map in two views (transparent surface) with model autobuilt by Modelangelo (ribbons, spectral coloring)

**Figure S5:**
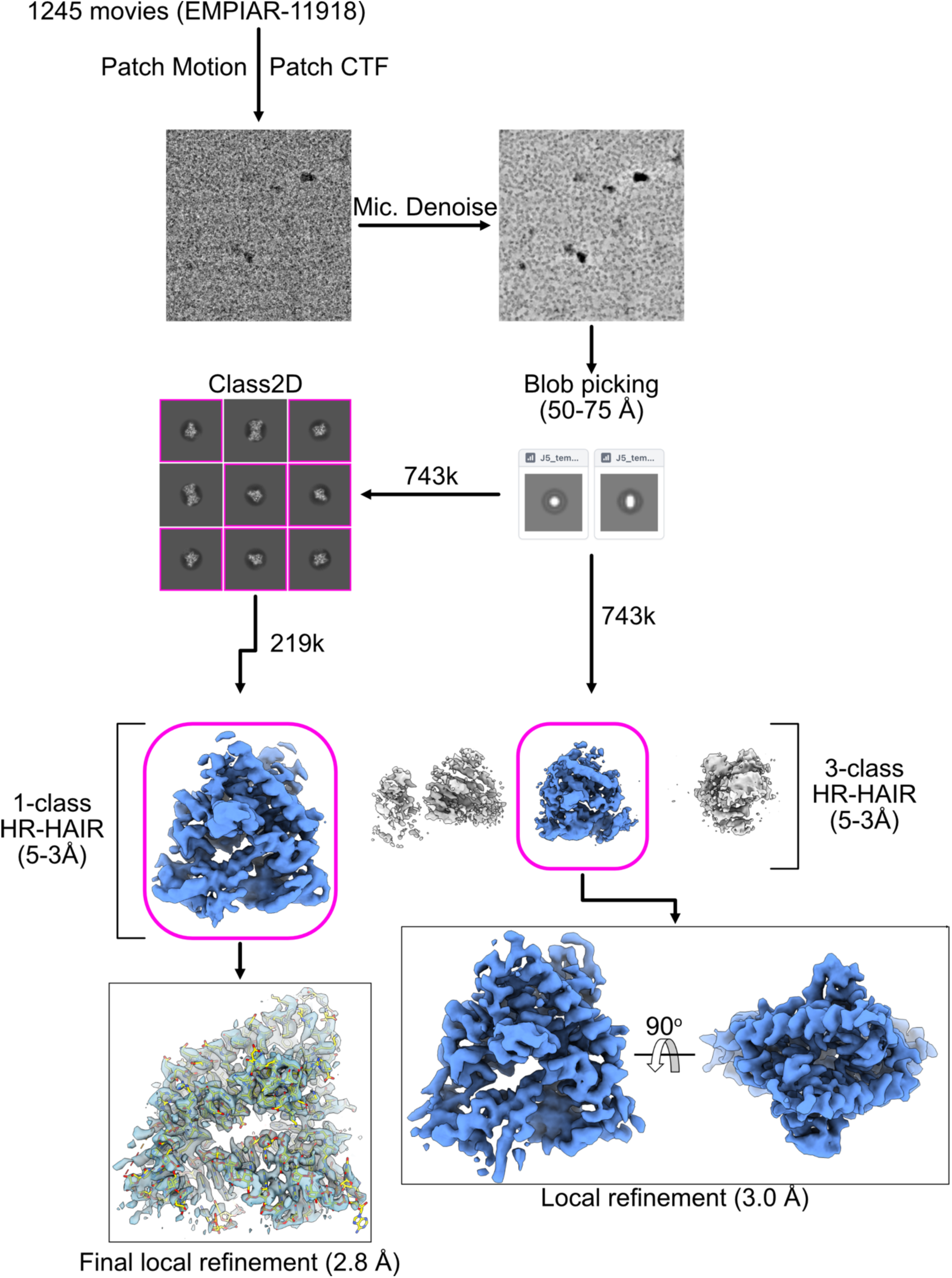
Aca2-RNA image processing schematic. Image processing workflow overview for Aca2-RNA; see Methods for details.

**Figure S6:**
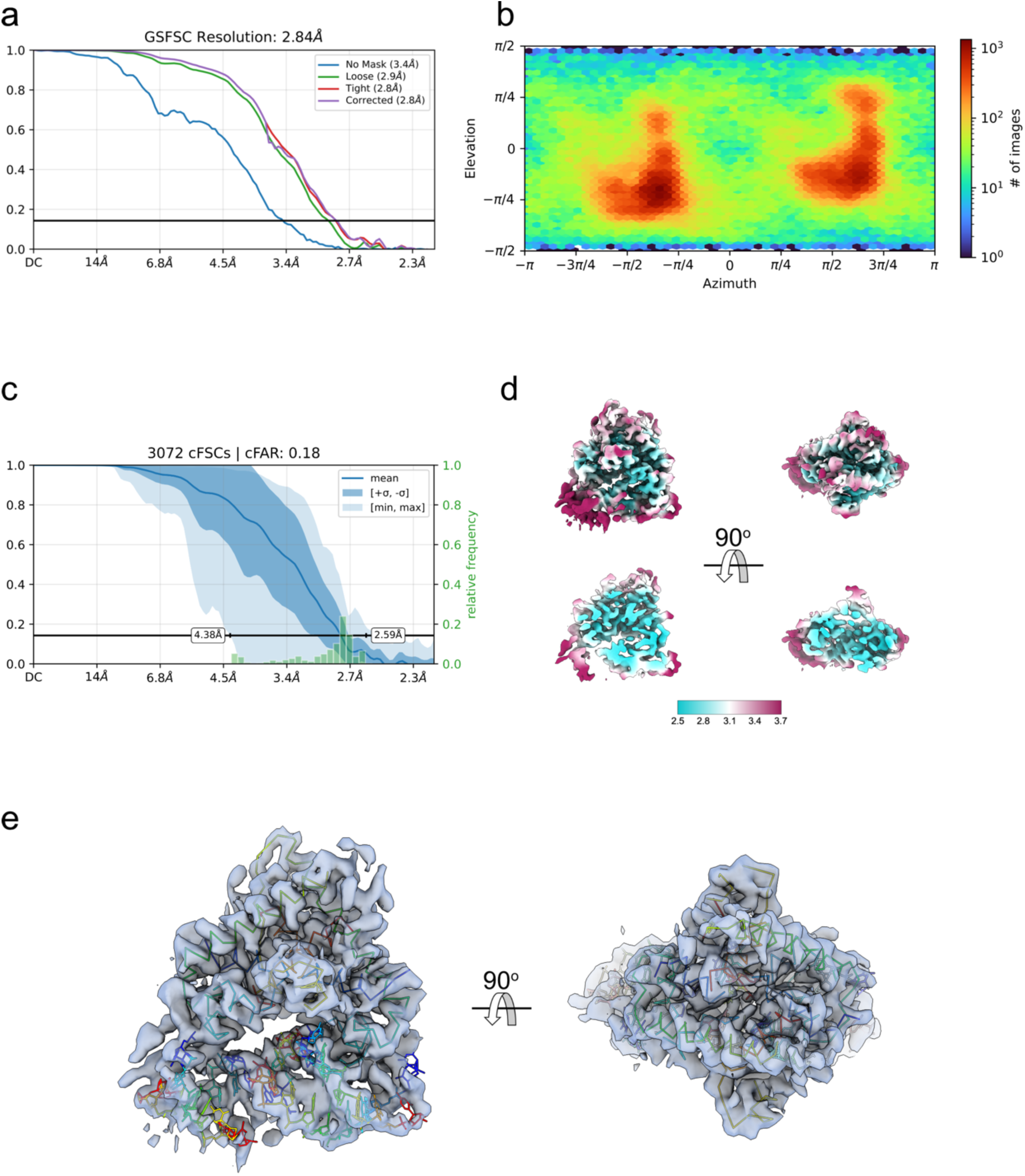
Aca2-RNA FSC curves, orientation distribution and local resolution. a) FSC curves after local refinement. b) Particle orientation distribution. c) Conical FSC distribution and conical FSC area ratio. d) Two views of the unsharpened map colored by local resolution, calculated by the local resolution module of CryoSPARC, with an FSC=0.5 threshold. Upper panels show an overall view of the map; below, the same view, sliced down the middle in the plane of the page. e) Unsharpened map in two views (transparent surface) with model autobuilt by Modelangelo (CA trace for protein, all atoms for nucleic acid, spectral coloring)

**Figure S7:**
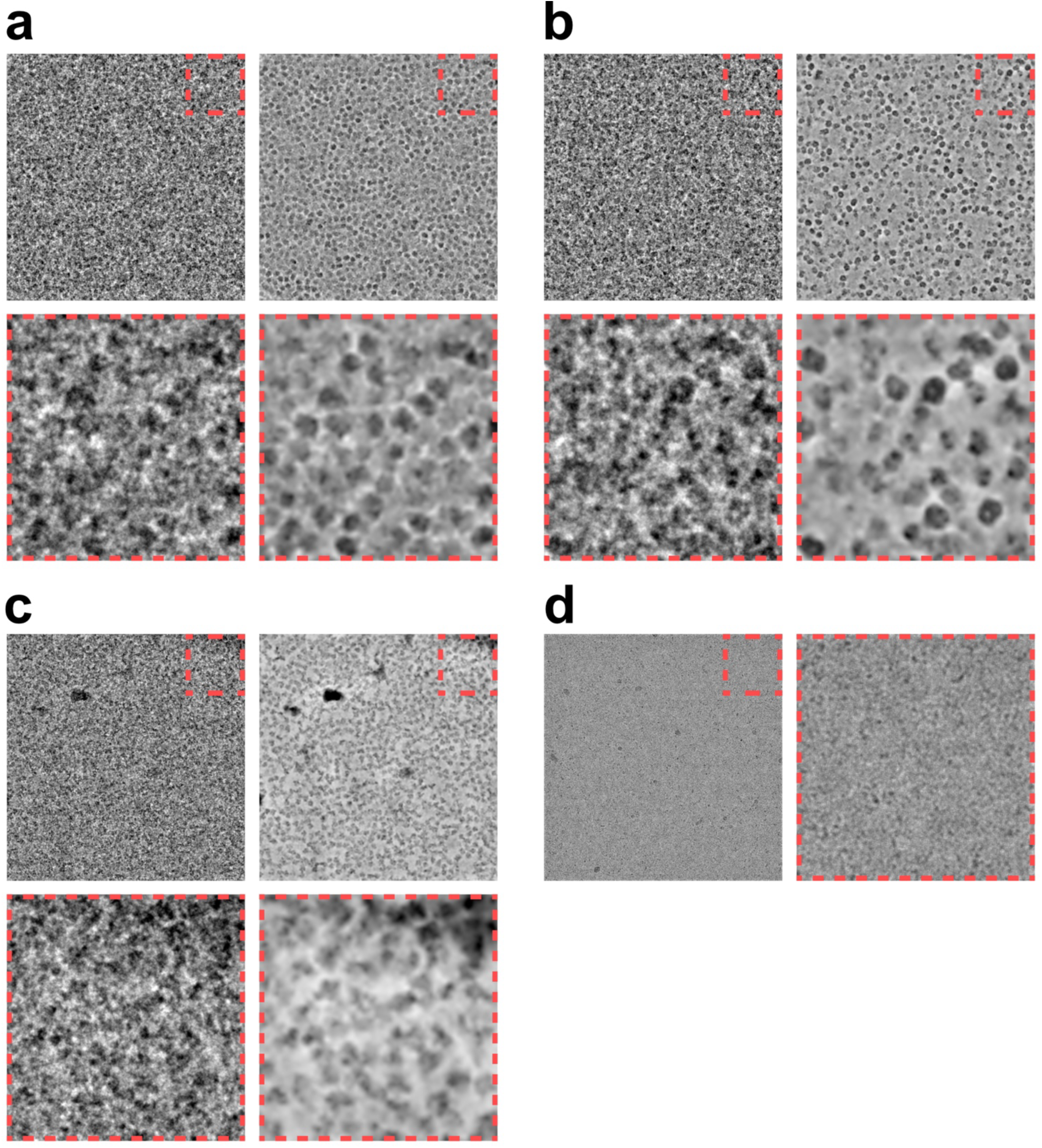
Comparison of original and denoised micrographs. a, b, c, and d correspond to iPKAc, Hb-dimer, Aca2-RNA and calmodulin, respectively. For a, b, and c, the left panels are not denoised, and lowpass filtered to 10 Å. The righthand panels show the denoised micrograph generated using CryoSPARC Micrograph Denoiser with no additional lowpass filter. The lower panels show a magnified view of the upper right 1/16^th^ of the micrograph, indicated with a red outline. Micrographs in a, b and c are at 0.7 µm defocus; the micrograph in d is at 1.5 µm defocus. For a, b and, d, the entire micrograph is shown, whereas for c, the Aca2-RNA micrograph (K3 detector) is cropped horizontally to match the dimensions of the other micrographs (K2 detectors). In d, a single micrograph of calmodulin denoised with Topaz denoise is displayed.

**Movie S1:**
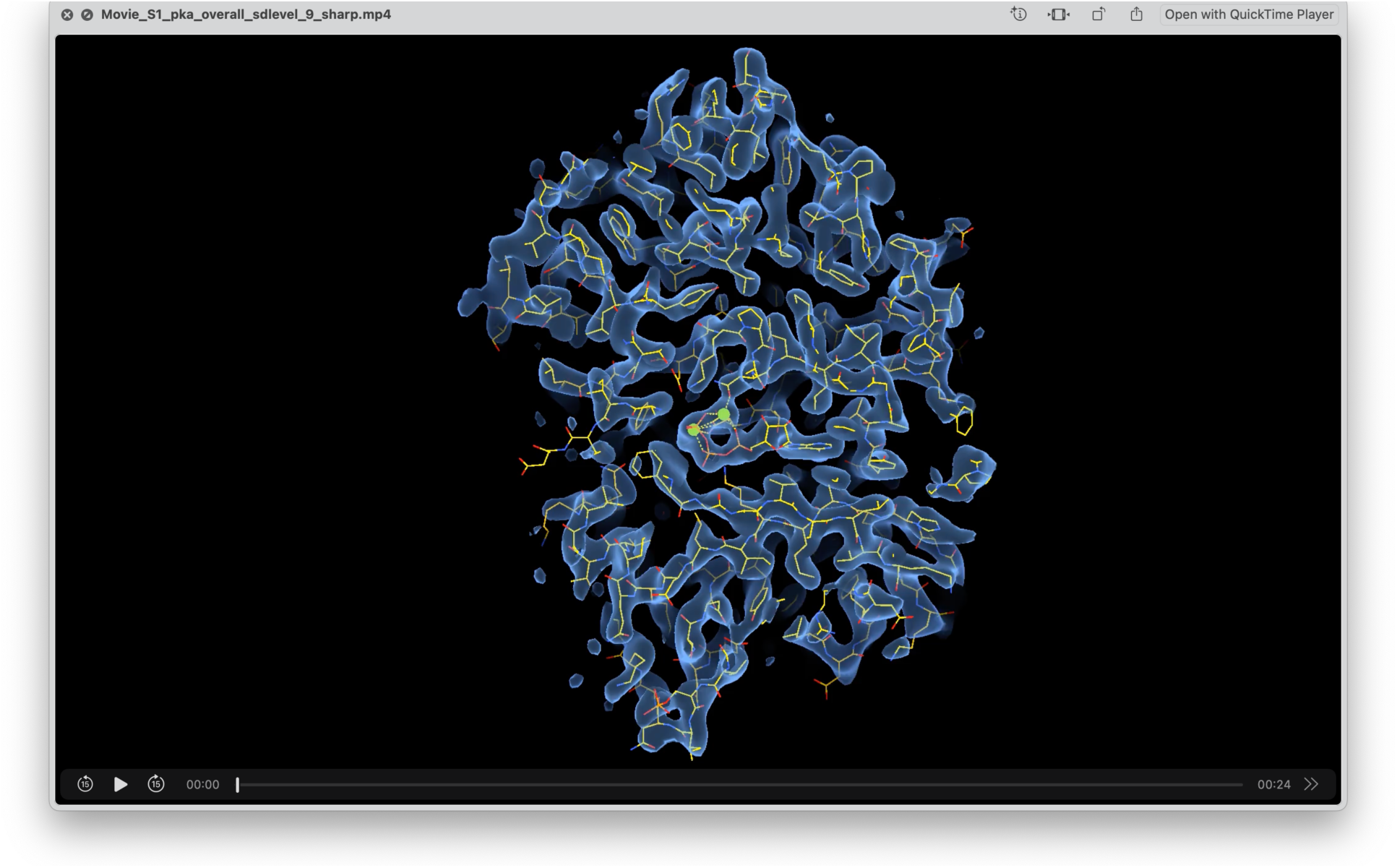
overall map/model fit for iPKAc. Map is represented as a transparent surface, contoured at an sdlevel of 9 in UCSF ChimeraX.

**Movie S2:**
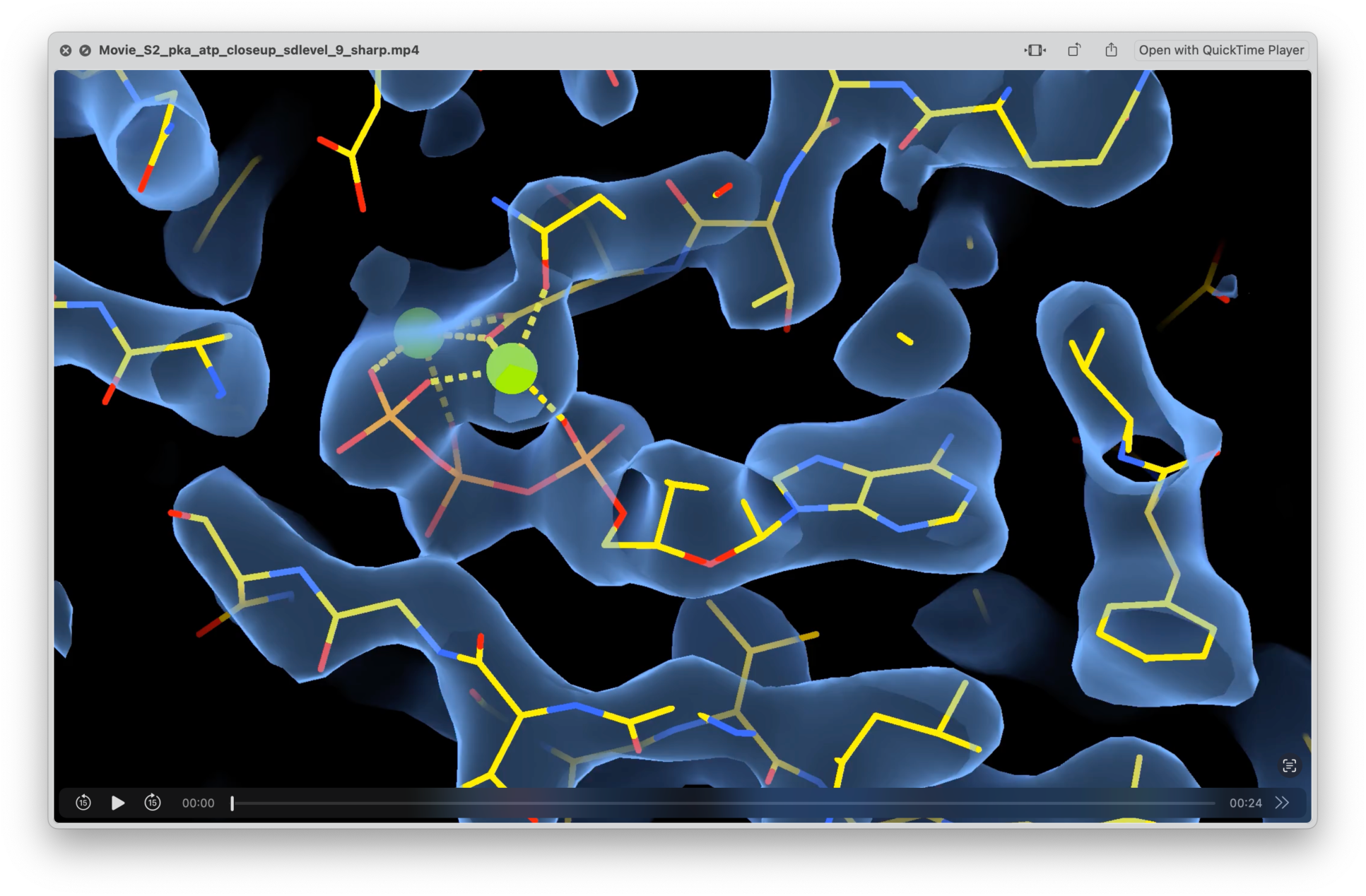
Closeup of Mg-ATP in iPKAc structure. Sharpened map is represented as a transparent surface, and contoured at an sdlevel of 9 in UCSF ChimeraX.

**Movie S3:**
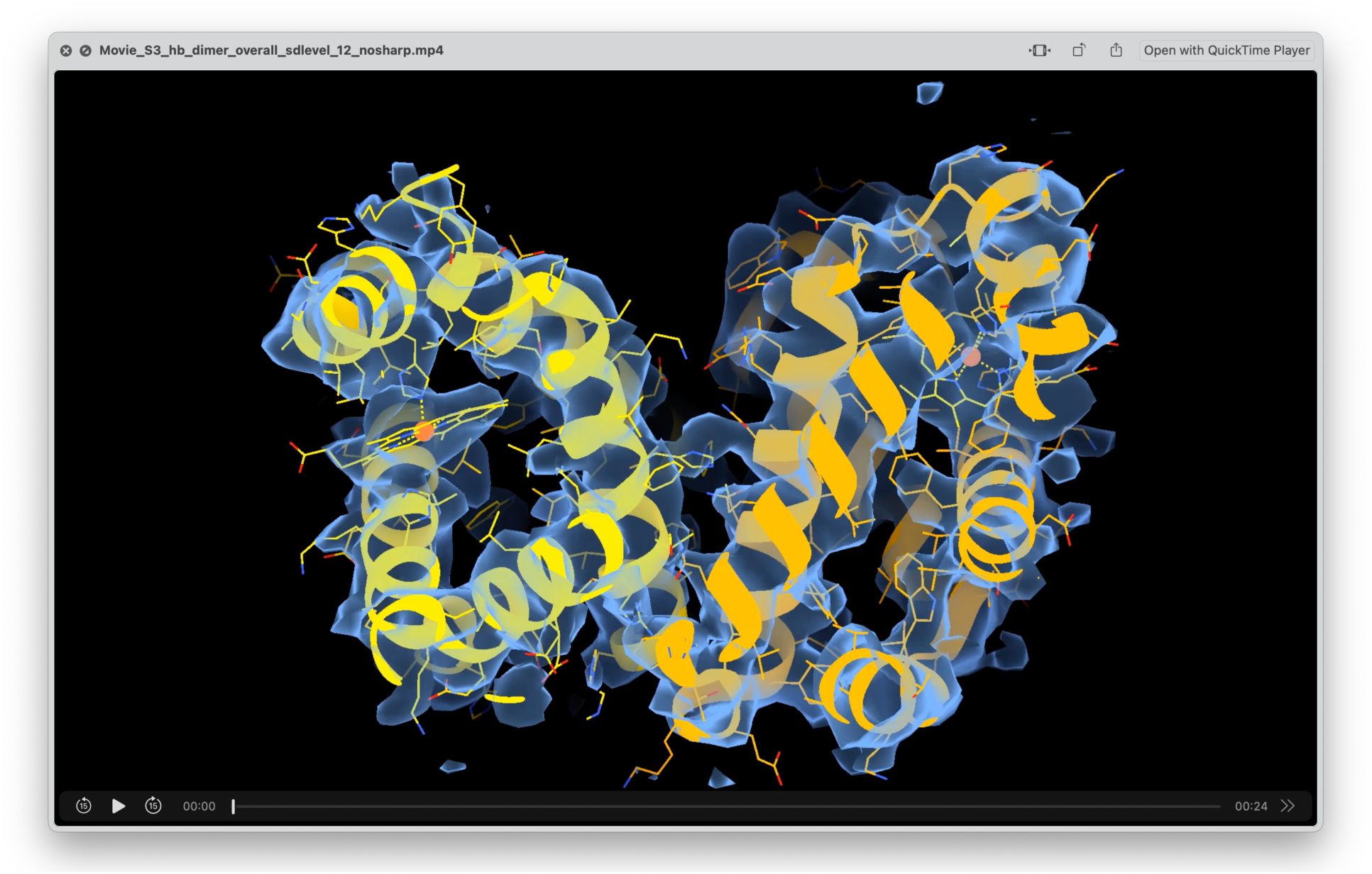
Overall map/model fit for Hb-dimer. Unsharpened map is represented as a transparent surface, and contoured at an sdlevel of 12 in UCSF ChimeraX.

**Movie S4:**
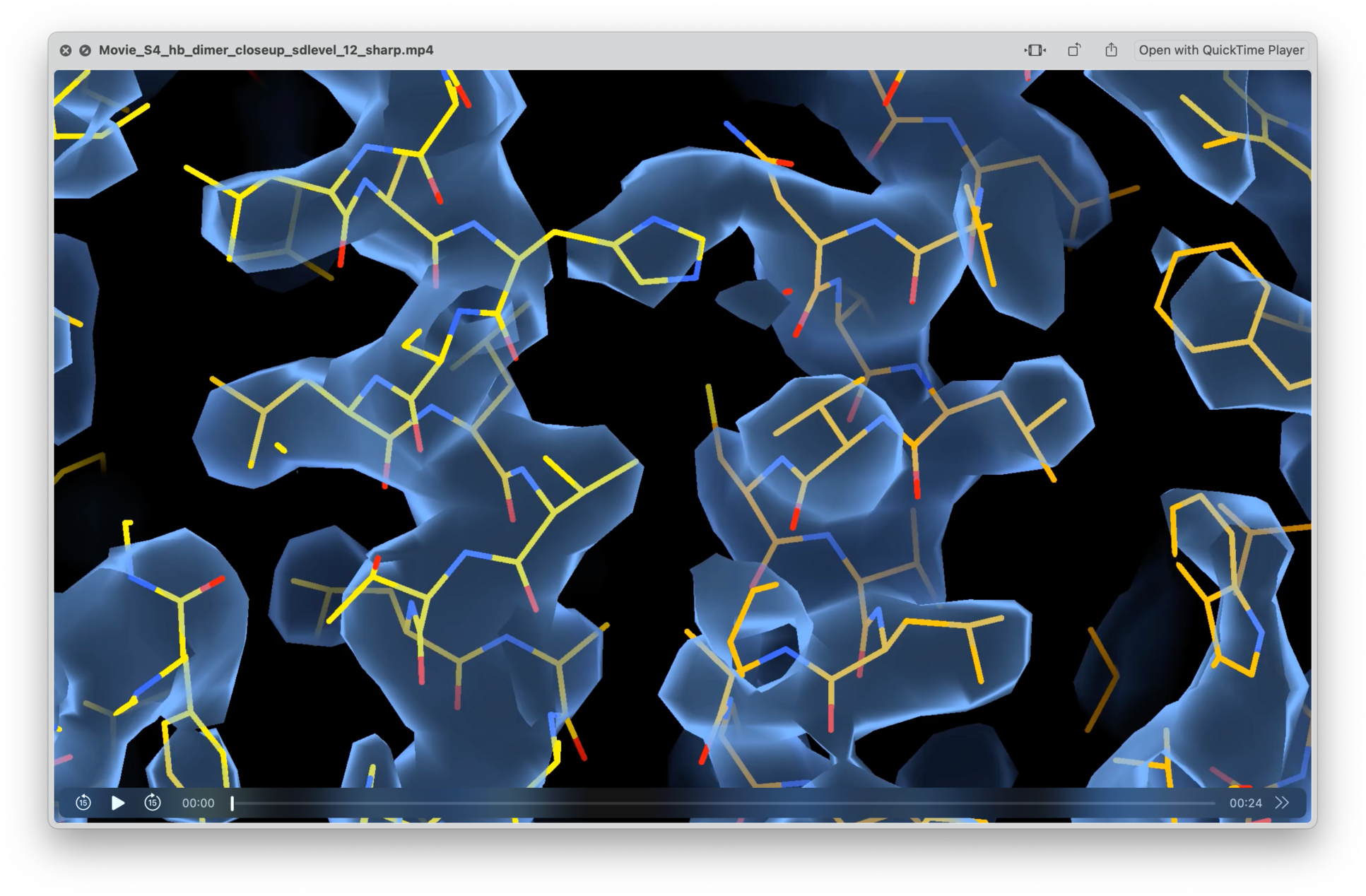
Closeup of the dimer interface of the Hb-dimer. Sharpened map is represented as a transparent surface, contoured at an sdlevel of 12 in UCSF ChimeraX.

**Movie S5:**
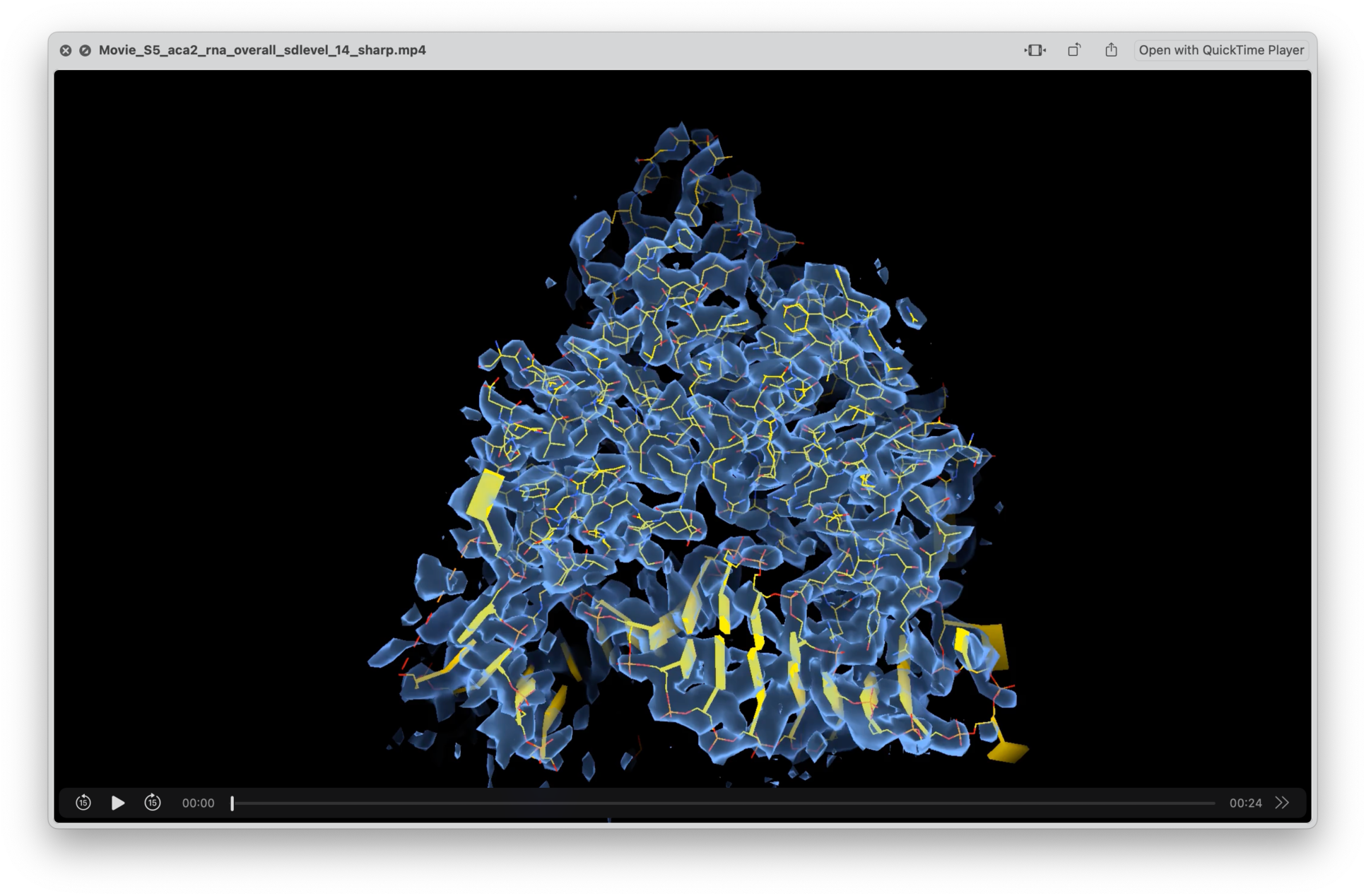
Aca2-RNA overall map/model fit. Map is represented as a transparent surface, and contoured at an sdlevel of 14 in UCSF ChimeraX.

**Movie S6:**
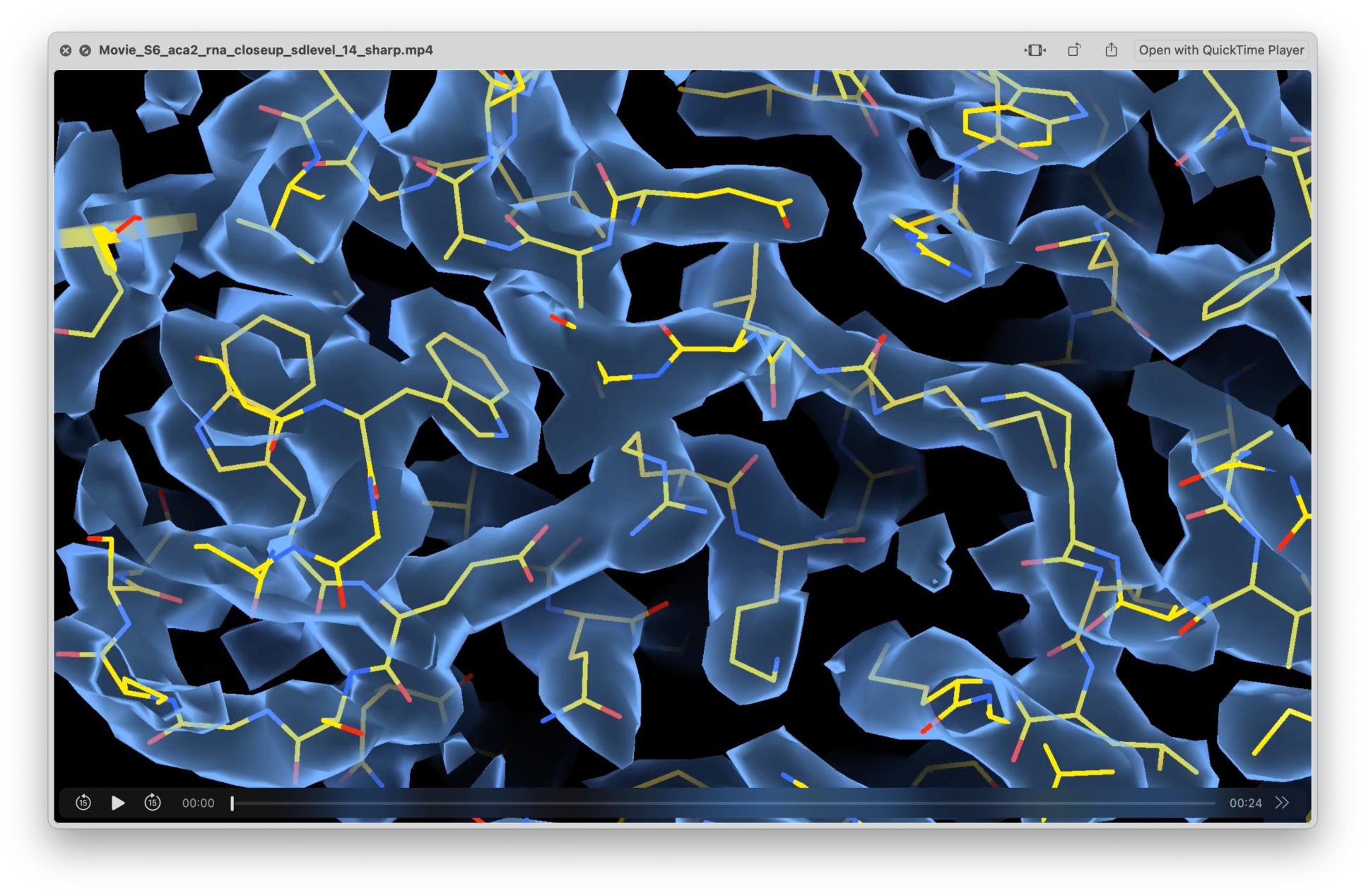
Closeup of Aca2-RNA map/model fit. Map is represented as a transparent surface and contoured at an sdlevel of 14 in UCSF ChimeraX.

**Movie S7:**
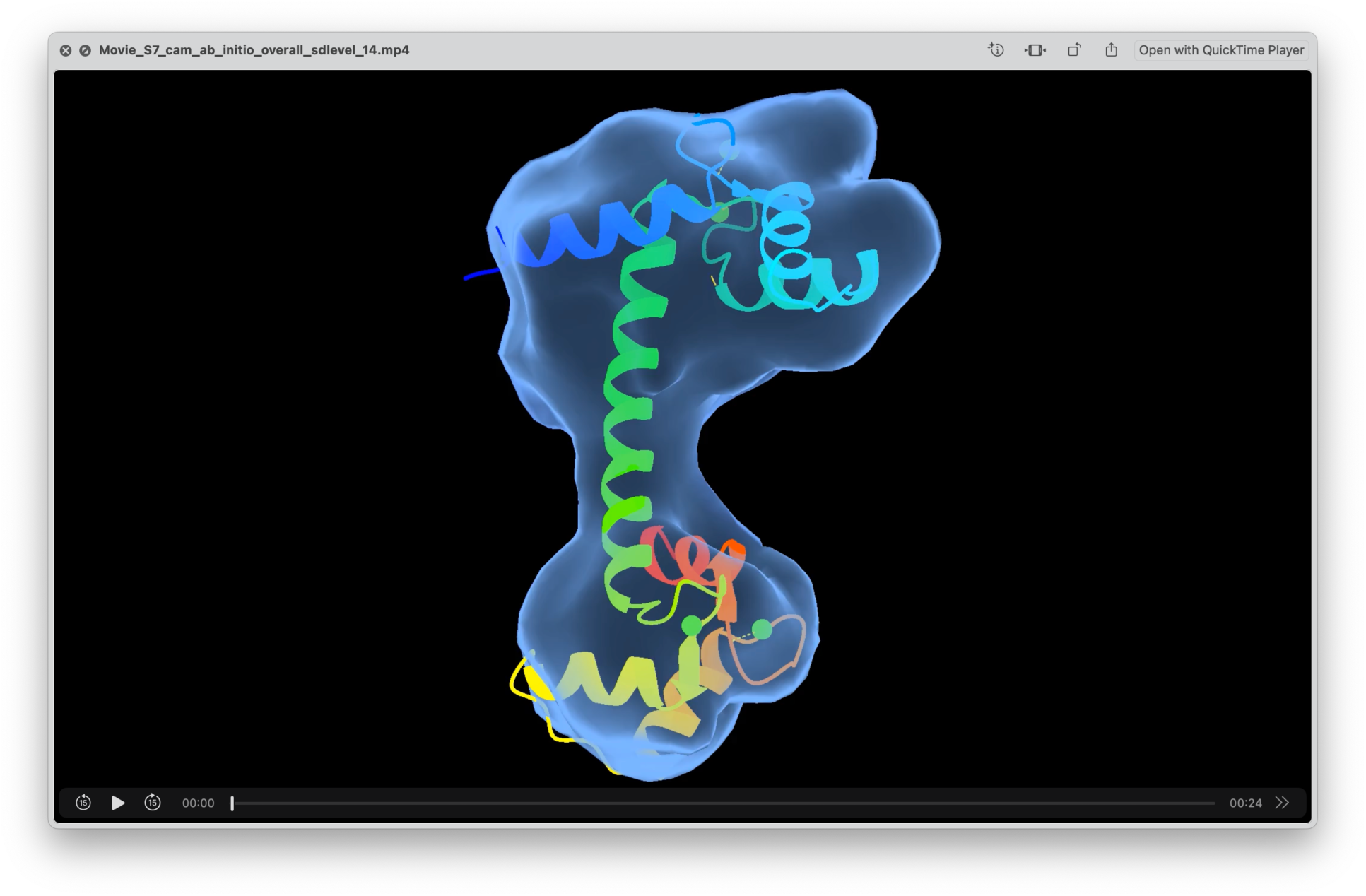
Calmodulin map/model fit. Map is shown as a transparent surface and contoured at an sdlevel of 14 in UCSF ChimeraX.

